# Strain population structure varies widely across bacterial species and predicts strain colonization in unrelated individuals

**DOI:** 10.1101/2020.10.17.343640

**Authors:** Jeremiah J. Faith, Alice Chen-Liaw, Varun Aggarwala, Nadeem O. Kaakoush, Thomas J. Borody, Hazel Mitchell, Michael A. Kamm, Sudarshan Paramsothy, Evan S. Snitkin, Ilaria Mogno

**Author notes:** Corresponding author (J.J.F.).

## Abstract

The population structure of strains within a bacterial species is poorly defined, despite the functional importance of strain variation in the human gut microbiota on health. Here we analyzed >1000 sequenced bacterial strains from the fecal microbiota of 47 individuals from two countries and combined them with >150,000 bacterial genomes from NCBI to quantify the strain population size of different bacterial species, as well as the frequency of finding the same strain colonized in unrelated individuals who had no opportunities for direct microbial strain transmission. Strain population sizes ranged from tens to over one-hundred thousand per species. Prevalent strains in common gut microbiota species with small population sizes were the most likely to be harbored in two or more unrelated individuals. The finite strain population size of certain species creates the opportunity to comprehensively sequence the entirety of these species’ prevalent strains and associate their presence in different individuals with health outcomes.

## Introduction

Although it was once unclear if bacterial species could be defined genomically^1,2^, the recent expansion of bacterial genomes, often with numerous isolates sequenced per species, has enabled genomic bacterial species definitions that empirically reflect or improve existing species names^3,4^. These species boundaries can be detected in the SNPs of conserved genes (i.e., the average nucleotide identity; ANI)^3–6^ and in the large differences in genome overlap (e.g., by pairwise genome alignment) or gene flow discontinuities driven by the strong bias of horizontal gene transfer within a species rather than across species boundaries^3,7–9^. As in microbial pathogenesis^10,11^, the functional impact of the microbiome is dependent on strain-level variation within a species^12–18^, which has driven computational advances to track strains^19–22^, cluster strains^23^, measure strain stability^7,21,24^, and analyze strain variation^25,26^. Strain-focused algorithms for both the commensal microbiome and infectious disease research have also begun to inform genomic boundaries for bacterial strains^7,21,22^. Despite the importance of strain-variation, we still lack a broad understanding of the general principles of strain population structure, such as the number of strains in each bacterial species, the stability of these strains^27^, the prevalence of each strain within a species in human and non-human reservoirs, and the fitness differences and environmental changes that drive alterations in strain prevalence^27,28^.

The study of bacterial pathogens provided the first genomics-based evidence of strain transmission and prevalence across human populations. The strong phenotype induced by both frank bacterial pathogens and colonizing opportunistic bacterial pathogens (COP)^29^ has facilitated the isolation and genome sequencing of numerous pathogenic isolates, which has demonstrated that pathogens within a given outbreak typically represent one or a small number of genomically-distinct lineages^10,30–32^. These results demonstrate that the same strain of bacteria can be harbored in multiple unrelated individuals. For many frank bacterial pathogens, this sharing is not through direct human-to-human transfer, but rather the same bacterial strain is colonized in unrelated individuals through the consumption of the same contaminated source (typically food or water).

For COP including *Clostridioides difficile* and extraintestinal pathogenic *Escherichia coli* (ExPEC), the colonization of the same strain in unrelated individuals can be both environment-to-human (e.g., shared occupation of a health care facility with insufficiently sterilized equipment) or human-to-human, as these organisms can stably and asymptomatically colonize the human gut and act as the reservoir for the recurrent reinfection of the target site of pathogenesis such as in urinary tract infections (UTI)^29^. Sequencing and isolation efforts for COPs initially focused on outbreak tracking within hospital intensive care units^30^. They have subsequently demonstrated that multiple COP strains are often asymptomatically maintained in long-term care facilities in a complex network of direct and indirect strain sharing^33^. COP strains are often multidrug resistant organisms (MDRO) whose antibiotic resistance may in part influence their prevalence in the human population which in turn increases the human risk for COPs. Broader sequencing efforts of MDRO COPs demonstrate that most strains cannot be explained by acquisition at healthcare facilities^32,34,35^ and are likely acquired elsewhere and stably maintained at various prevalence in the healthy human population.

Understanding bacterial population structure and strain prevalence, beyond the narrow lens of the hospital environment, could provide novel tools to quantify the association of microbial strains with both infectious and complex human disease, as well as new routes to limit human disease. Early broad explorations of the gut microbiota have demonstrated contexts of enriched strain sharing for commensal microbes including the shared hospital environment for infants^22^, fecal microbiota transplantation (FMT)^19,21^, and early life co-habitation between family members^7,22^. Although selective pressures likely differ in non-MDRO organisms, the prevalence of a bacterial strain in the broader human microbiota population, beyond enriched scenarios of direct transfer like co-habitation and FMT, is a reflection of a strain’s fitness that includes both the transmissibility of the organism and its stability in the host^13^.

Here we use a sequenced collection of 2359 bacterial isolates representing 1255 strains that were isolated from the fecal microbiota of 47 individuals from USA and Australia, to study principles of strain population size and their implications for strain-prevalence in unrelated individuals. Strain population size varies dramatically across species with some species being represented by tens of strains and others represented by hundreds of thousands. Prevalent bacterial strains from species with small strain population sizes are far more likely to be colonized in two unrelated individuals than strains from species with large strain population sizes. The finite number of bacterial strains within each species creates the potential to track them and their genetic loci across individuals to identify those associated with short- and long-term health outcomes for both COP pathogenesis and complex disease.

## Results

### Defining a bacterial strain as a pairwise genome kmer overlap of 0.98 or greater

To better understand the strain population structure of species resident in the human microbiome, we generated a dataset combining 156,403 bacterial genomes from NCBI with 2359 newly sequenced bacterial isolates (hereafter referred to as LOCAL) from 257 species isolated from 47 individuals across two countries (USA^14,21,36^ and Australia^37,38^). We used a k-mer hash-based approach to efficiently calculate the genome overlap between all pairs of bacterial genomes from the same species as the proportion of shared k-mers between the genomes. As in our previous work^7^, we find these species-level genome comparisons are dominated by highly similar (>0.98 kmer overlap) genomes when comparing multiple isolates of the same species from a single individual at a single timepoint (Fig. 1A.i) – reflecting the situation where multiple isolates of the same strain are captured and sequenced from the same stool sample. Also similar to our prior work, we find that pairwise comparisons of isolates from the same species that were isolated from one individual at different time points are also dominated by kmer overlaps of >0.98 (Fig. 1A.ii), as these strains are stably maintained over time in each individual and re-isolated at a second time point^7,39^. Given this strong empirical observation of the kmer overlap of >0.98 between genomes of the same species isolated from an individual, we will use 0.98 as the threshold for defining a bacterial strain for the remaining comparisons across individuals. Performing these analyses with the other popular pairwise genome comparison method of Average Nucleotide Identity (ANI)^5^ yields similar results. However, we find that kmer overlap, which captures both SNPs and gene flow discontinuities^9^, provides improved resolution with less signal saturation between very similar species (Fig. S1A) and better identifies isolates sequenced from the same individual (Fig. S1B).

**Figure 1.**
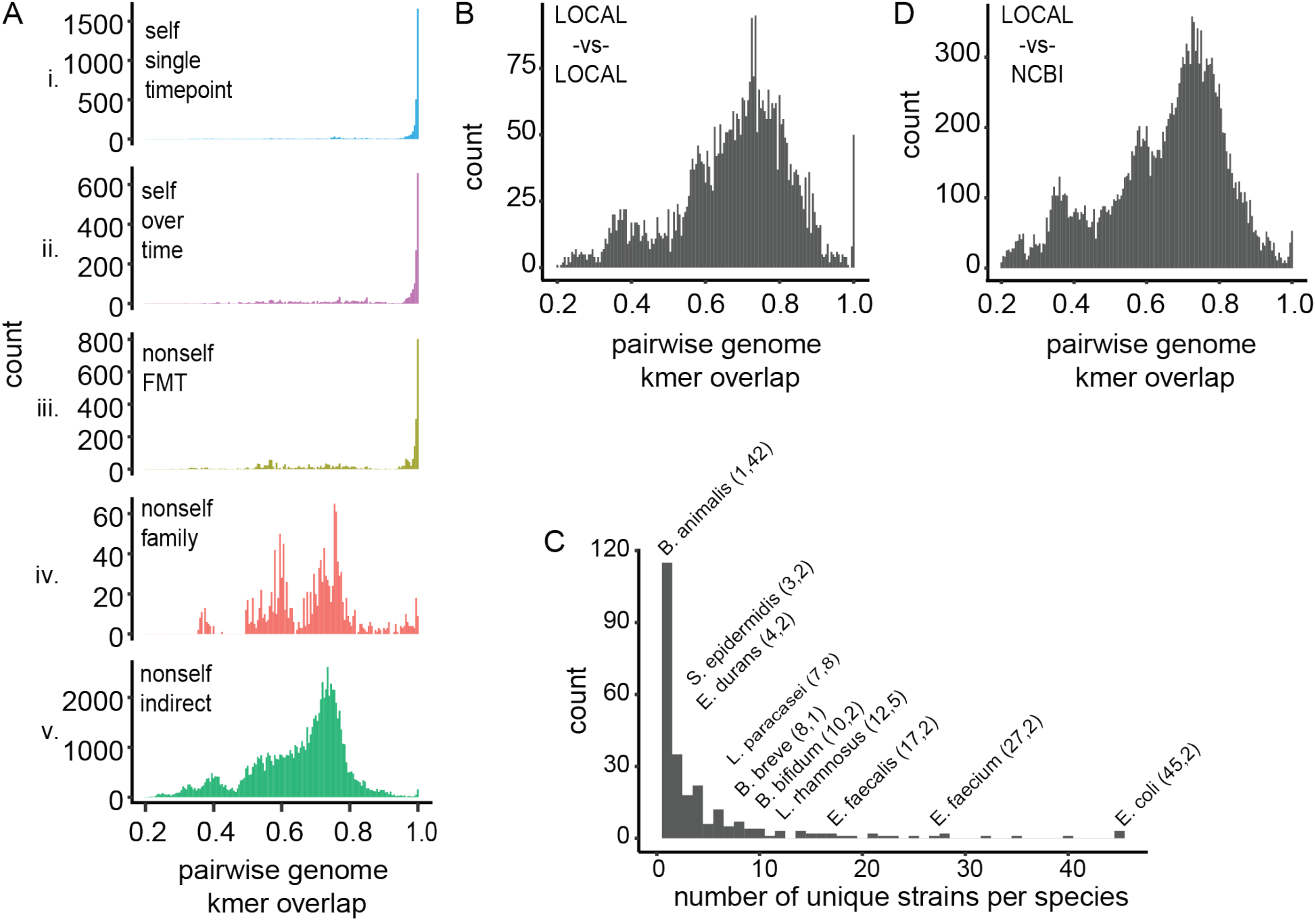
Highly similar bacterial species are enriched in the context of cohabitation and transmission but not absent in all unrelated individuals with no direct contact. (**A**) The shared kmer content was calculated for all pairwise combinations of species between (i) individuals’ own microbes from a single sample, (ii) individuals’ own microbes from two different timepoint, (iii) FMT donors and their recipients, (iv) members of the same family, (v) two unrelated individuals with no opportunities for direct microbial transfer between them. (**B**) A randomly subsampled set of all pairwise species kmer overlap between genomes from 47 different individuals reveals a peak at kmer distance >0.98 even when eliminating strains assumed to be shared by direct transmission (FMT) or cohabitation (family). (**C**) The strains indirectly shared in LOCAL were from nine different bacterial species with varying numbers of strains in our genome set. For species labels on pane C, the first integer is the number of unique strains for a given species in LOCAL, while the second integer is the number of pairwise observations of the same strain in two unrelated individuals with no direct transfer event. (**D**) A similar high kmer overlap >0.98 peak was observed between LOCAL genomes and NCBI genomes.

### Bacterial strains can colonize different individuals by direct transfer

Although it is clear that pairs of individuals do not typically have large overlaps in their microbiome strain composition, cohabitation and fecal microbiota transplantation provide two possibilities for direct strain transmission between individuals to perhaps increase their strain composition overlap. Comparing the kmer overlap between genomes from the same species isolated from fecal microbiota transplant (FMT) donors and their recipients treated for recurrent *Clostridioides difficile* (rCDI)^40,41^, kmer overlaps >0.98 again dominate, with perhaps slightly more kmer overlaps <0.98 demonstrating the likely acquisition of strains from non-donor (environmental) sources after the transplant (Fig. 1A.iii). These results are in line with our recent observation that the recipient microbiota post-transplant is composed of 80% donor strains, 10% recipient strains (i.e., those colonizing the recipient prior to transplant), and 10% environmentally acquired strains^21^. The second category for an increased chance for direct transmission of bacterial strains is between family members. Although this bacterial transmission is less purposeful than the large microbial biomass transferred in FMT, the cohabitation of individuals provides numerous opportunities for microbial transfer, particularly in early life as the gut microbiota is established^13^. These familial pairwise genome comparisons from isolates of the same species again contained numerous kmer overlaps of >0.98 suggesting familial transfer of bacterial strains as demonstrated in prior studies (Fig. 1A.iv)^7,42^. Notably, these strain-level kmer overlaps were a minority suggesting direct transmission within families is only a fraction of that experienced via FMT.

### Bacterial isolates with >0.98 genome kmer overlap can be found in pairs of individuals without direct transfer

Although it is clear that factors such as direct transfer of strains via FMT can facilitate strain sharing of commensal bacteria between unrelated individuals and that there is substantial species overlap between individuals, the number of strains in a bacterial species could be sufficiently high or the mutation and recombination rate could be so rapid that it is unlikely to find the same commensal bacterial strains colonized in unrelated individuals inhabiting distal sites of the Earth. We performed pairwise genome kmer overlap comparisons between genomes of the same species from all remaining isolates in our dataset consisting of unrelated individuals where no direct transmission of strains was likely (i.e., no FMT and the individuals are unlikely to have ever had direct contact). We find that the vast majority of comparisons are <0.98 kmer overlap. However, there is a slight, but notable, peak in kmer overlap at an overlap of 0.98 and greater (Fig. 1A.v). This peak could indicate that the population sizes of some common bacterial species are sufficiently small that with our cohort of only 47 individuals we are finding unrelated individuals harboring the same bacterial strain or that convergent evolution within a species pangenome repeatedly drives to a highly similar state.

### Bacterial strains with >0.98 genome kmer overlap are found for frank pathogens and COP pathogens

For comparison with frequent colonization of the same strain in unrelated individuals in complex disease, we calculated the kmer overlap between environmental and human isolates from a spinach outbreak and a “Taco John” outbreak of frank pathogen *Escherichia coli* O157:H7^31^. The mean+/-std kmer distance of genomes from within each outbreak was 0.982±0.018 and 0.985±0.017 for spinach and “Taco John” respectively, while the kmer distance of genomes compared between the two outbreaks was 0.955±0.189. Applying the same method to MDRO COP in the context of carbapenem-resistant *Klebsiella pneumonia* outbreaks in Beijing Tongren hospital^30^ and Shanghai Huashan hospital^43^, we found all individuals in the Beijing Tongren outbreak were infected by the same strain (0.996±0.004) while individuals in the Shanghai Huashan outbreak had one of four different strains (0.994±0.004). Similar to the frank pathogens, the kmer distance between independent outbreaks was 0.951±0.020. These results demonstrate that the kmer distance of strains shared in the commensal microbiota can be similar to lineages in pathogen outbreaks.

### Colonization of the same strain in two unrelated individuals with no direct transfer is more prominent when species comparisons are more evenly represented

To further probe the extent to which unrelated individuals might harbor the same bacterial strain, all remaining analyses are focused on pairwise comparisons of LOCAL isolates from the same species between individuals with no direct transfer opportunities (e.g., as in Fig. 1A.v). A caveat of our comparisons in Fig. 1A is that our LOCAL dataset has an uneven number of representatives from each species reflecting both their prevalence in the human population and their ease of bacterial culture. Therefore, in performing all possible pairwise comparisons of isolates in each species, the more common species in the LOCAL dataset will have far more comparisons than those that are rare. To better reflect the kmer overlap distribution across species, we randomly subsampled the pairwise kmer comparisons for each species to have at most the same number of comparisons as the species whose prevalence was the upper quartile LOCAL dataset (Fig. 1B). As expected from prior work^3,7^, organisms from the same species have a characteristic genome overlap where the most common overlap is around 0.70 with increasingly similar kmer overlaps diminishing sharply from kmer overlaps of 0.70 to 0.97 (Fig. 1B). Intriguingly when we use this subsampling approach to include more proportional representation of less frequent species, this decay is followed by a sharp peak of genome kmer overlaps of 0.98 to nearly 1.00. Focusing on all genome kmer overlaps >0.98 grouped by species further reveals that this peak is heavily weighted towards very high genome kmer overlaps of >0.995 with few pairwise comparisons near 0.98 (Fig. S1C). Given the natural decay of genome similarity after 0.70, it seems highly unlikely that this second higher peak occurred by chance. It likely reflects a small subset of strains colonized in multiple unrelated individuals who harbor the same strain but not through direct microbial transmission. These shared strains were found in species for which we have numerous distinct strain isolates and those species with as few as a single unique strain (Fig. 1C) suggesting this observation is not simply an artifact of sampling bias where more prevalent species have more genomes increasing the chance of finding two unrelated individuals with the same strain. Across the ~20,000 pairwise comparisons of LOCAL isolates from the same species between individuals with no direct transfer opportunities, only 0.35% were >0.98 kmer overlap, encompassing 4.67% of the 237 species and 1.35% of the 1255 strains. Amongst all pairwise comparisons of the 47 individuals in the cohort, the chance of a pair of individuals harboring at least one strain that is the same between them was 3.4%, and only one pair of individuals shared two strains that were the same – at the strain level human microbiomes are almost totally unique. While all of the bacterial isolations in the LOCAL dataset were performed in a single anaerobic chamber, these shared strains were often isolated from culture libraries generated years apart, mitigating the chance they represent contaminants.

### Colonization of the same strain in two unrelated individuals with no direct transfer is confirmed in public bacterial genome databases

The large number of publicly available bacterial genomes in NCBI provide an independent dataset to validate the enrichment of genomes with a pairwise kmer overlap of greater than 0.98 in the absence of direct strain transfers between pairs of individuals. While the composition of the strains in NCBI is likely biased towards commercially and medically relevant strains in some species, we can limit these biases to a large extent by comparing the LOCAL bacterial genome to those in NCBI. Although we have limited metadata on the NCBI bacterial genomes, it is highly improbable that the LOCAL strains were isolated from individuals that are first degree relatives or fecal transplant recipients of the individuals whose microbes are in the NCBI genome set. We calculated the kmer overlap of LOCAL bacterial genomes with NCBI bacterial genomes. As in our LOCAL comparisons, the number of representative genomes for each species is highly varied, and we randomly subsampled the pairwise kmer comparisons for each species to have at most the same number of comparisons as the species whose prevalence was the upper quartile in LOCAL dataset (Fig. 1D). We again found a kmer overlap spike between 0.98 and 1.00 suggesting the same bacterial strain is found between unrelated individuals in two independent datasets (i.e., within LOCAL and between LOCAL and NCBI) (Fig. 1D). As with the LOCAL pairwise comparisons, we also find for most species that the kmer overlaps >0.98 are heavily biased towards very high overlaps of >0.995 (Fig. S1D). *Enterococcus faecalis* and *Escherichia coli* are two notable exceptions to this trend, as their pairwise interactions look more like the true decay of the tail of a distribution rather than a second peak.

### Strain population sizes vary widely across bacterial species

If unrelated individuals are harboring the same strains of bacteria, it suggests that bacterial species have a finite number of strains (i.e., a population size) that are stably maintained and propagated in the human population. In both macro- and microecology, it is often impossible to exhaustively sample a population to determine its size (also known as total taxonomic richness). Two approaches are commonly used to infer population sizes from a subsample of its members. One of these approaches takes a subsample of the population (e.g., the set of strains in species *Bacteroides ovatus* in NCBI) and quantifies the frequency distribution of population members found once, twice, etc.. as f_1_, f_2_…, f_N_ (Fig. 2A). If the population is not exhaustively sampled, there exists an unobserved group f0 that has not yet been detected in the subsample, which can be inferred from the data (e.g., the number of unique *B. ovatus* strains that have not yet been isolated and sequenced)^44^. After inferring f_0_, the population size can then be calculated as the sum of the observed and unobserved community members with the assumption in our case that the number of strains within a species at any time (or within the timescale of human lifetimes) is finite. We applied the iChao algorithm of Chui and Chao^45^ that uses Hill statistics to estimate strain population sizes. To focus on the gut microbiome and species where this inference would be most robust, we calculated strain population sizes for the species in LOCAL that had at least 50 genome sequences in NCBI and that were found shared within LOCAL or between LOCAL and NCBI. Across these species, we inferred vastly different population sizes across a >9000-fold range with 19 as the smallest strain population estimated for *Bifidobacterium animalis* and 1.8×10^5^ estimated for *Escherichia coli* (Table 1).

**Figure 2.**
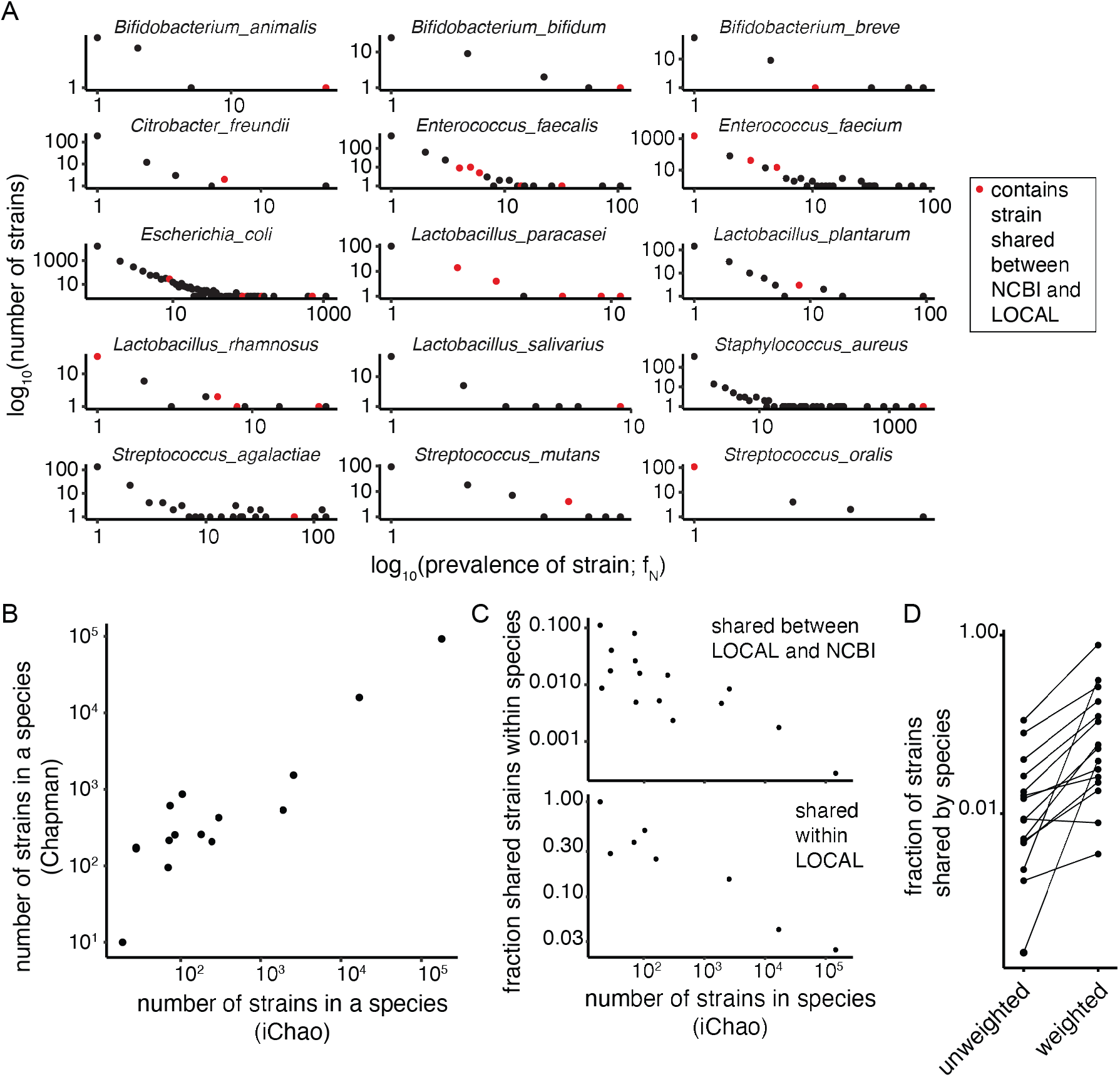
Strain population size varies by species and predicts the frequency of strain sharing between unrelated individuals with no direct microbial transmission. **(A)** Strains in NCBI are present at different frequencies with the largest number of strains present only a single time and a few prevalent strains that are much more highly represented in the species’ population sample of genomes available from NCBI. **(B)** Estimation of strain populations for bacterial species is highly correlated when using either a strain frequency approach (iChao) using only genomes from NCBI or a mark and recapture method (Chapman) that considers the proportion of LOCAL strains found in NCBI. (**C**) The strain population size of each species is highly predictive of the fraction of indirect strain sharing between LOCAL and NCBI (top pane) and indirect sharing within LOCAL (bottom pane). (**D**) The unweighted proportion of NCBI strains shared with LOCAL is the proportion of NCBI strains shared with LOCAL weighted by strain prevalence.

**Table 1.**
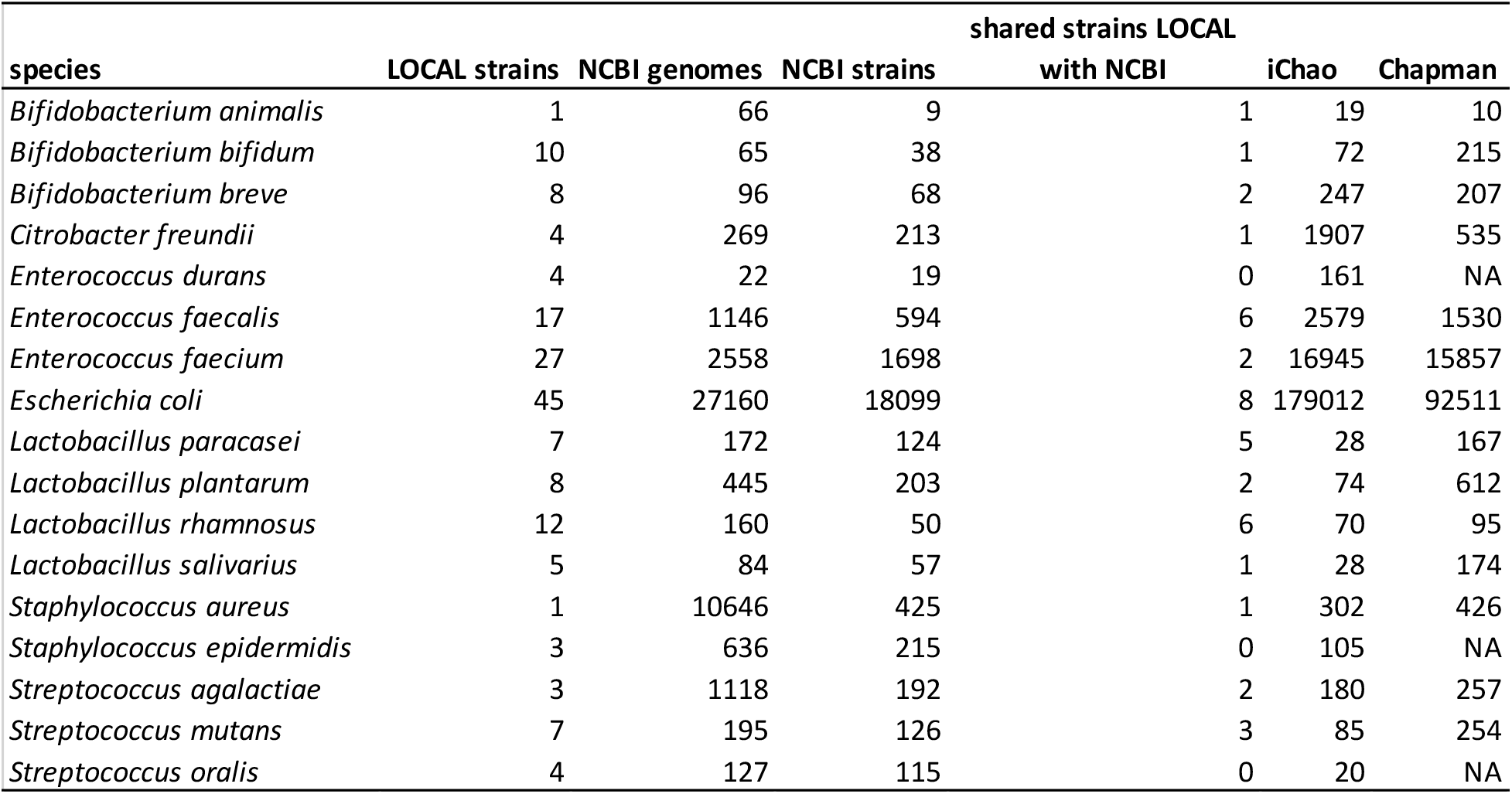
Strain population sizes for bacterial species.

Mark and recapture methods provide an alternative method to estimate population size by using two consecutive subsamples of a population. In the first sampling, the “captured” members are marked and released back into the population. In the second sampling, one will observe some new members and potentially some “recaptured” members that were marked in the previous round. Population size can be estimated from the number of members collected in each of the two subsamplings and the proportion of marked and unmarked community members resampling (e.g., using the Chapman algorithm^46^). To apply this alternative method of calculating population size, we assume the genomes in NCBI represent the initial sampling and the LOCAL genomes represent the resampling. Like the frequency distribution approach above, the mark and recapture approach estimated vastly different population sizes over a >9000-fold range with 10 as the smallest strain population estimated for *Bifidobacterium animalis* and 9.2×10^4^ for *Escherichia coli* (Table 1). The log of the strain population sizes for each species estimated with these two different approaches were highly correlated (r=0.91; p=2.9×10^-6^; Fig. 2B) suggesting they roughly approximate the true strain population size of each species.

Given the large variation in strain population size across species, our probability of observing indirect strain sharing, within a species between two individuals, will be influenced by the population size for that species. As expected, we find a significant negative correlation between the log proportion of strains shared within a species in LOCAL and the log population size for the species (r = −0.95; p=2.8×10^-4^; Fig. 2C top plot) as well as between log proportion of strains shared between LOCAL and NCBI for a given species (r = −0.81; p=2.4×10^-4^; Fig. 2C bottom plot). For example, *B. animalis* had a population size of 19 estimated from 66 genomes from 9 unique strains in NCBI. In LOCAL, *B. animalis* had single unique strain that colonized seven different individuals, five individuals in the USA and two individuals in Australia.

Just as more favorable genetic alleles expand in the human population, we would expect the frequency distribution of strains within a bacterial species to be uneven and in proportion to the fitness of each strain with the most transmissible and stable strains dominating the species. These more frequent strains within a species would similarly have an increased chance of being found in two unrelated individuals. To test this hypothesis, for each species shared between LOCAL and NCBI we compared the unweighted proportion of the strain within the species in NCBI to the weighted prevalence of the strain in the species in NCBI, where the weighted prevalence is defined by the frequency of the strain in NCBI. For example, we found one of the nine *B. animalis* strains in NCBI was shared with LOCAL (11%). However, since this shared strain was also the most prevalent *B. animalis* strain in NCBI (51 out of 66 genomes), it represented 77% of the *B. animalis* genomes in NCBI. It was similarly the case for all but one (*S. oralis)* of the 14 shared strains between LOCAL and NCBI that the shared strains represented more prevalent strains within a given species (p=0.026; paired t-test; Fig. 2D). This bias towards indirect sharing of prevalent strains can also be seen in Fig. 2A where the strain frequency in NCBI is highlighted in red for strains that are shared between NCBI and LOCAL. The red points are skewed towards the right showing that the more prevalent strains were more likely to be found indirectly shared between the two datasets.

## Discussion

Overall, we have identified numerous instances of the same bacterial strain harbored in two different individuals without a direct microbial transmission event. These results suggest the number of strains in at least some bacterial species can be finite and stably maintained in the human population where they colonize unrelated individuals across the world^47^. Predictably, common species with smaller population sizes were more likely to be found shared between individuals. Finally, strains were unevenly distributed within each species with presumably more fit strains being more prevalent in the population and more likely to be found in unrelated individuals.

Although genetic diversity is generated in bacteria through mutation and horizontal gene transfer and the strains within a community will drift as one or more stable substrains over time^39^, for many species the genetic boundary of 98% genome similarity appears to be retained at least on human health relevant time scales. Here we observed this phenomenon in the incredibly similar kmer overlaps between the strains shared between LOCAL and NCBI. For example, one shared strain of *Lacticaseibacillus rhamnosus* has a kmer overlap of 0.9994 with the type strain isolated >30 years ago^48^ and a shared strain of *Ligilactobacillus salivarius* has a kmer overlap of 0.9988 with a strain isolated >60 years ago^49^.

Notably, the strains harbored in pairs of unrelated individuals in LOCAL or between LOCAL and NCBI were limited to only 16 total species out of the 237 in our cohort. Species with the smallest strain populations were often ones used in probiotics, suggesting their small population size might have resulted from direct human intervention limiting strain diversity and increasing the prevalence of certain strains. Other species with smaller populations of strains were microbes that are found more commonly in other habitats including skin origin (Staphylococci)^50^ and oral origin (Streptococci). These organisms were perhaps transient components of the fecal microbiota, from the individual’s microbiome, that were enriched by selective media used to culture each fecal microbiome. This result might suggest that average strain population sizes will differ for species enriched in different habitats. Of the four dominant phyla of the human gut microbiota (Firmicutes, Bacteroidetes, Actinobacteria, and Proteobacteria), only Bacteroidetes were never found to be indirectly shared in our analyses, perhaps because the number of currently sequenced strains for all species in this phylum in both NCBI and LOCAL is low. Amongst the tested species in our analyses, only 4% and 13% of those with <10 unique strains and 10-100 unique strains respectively in NCBI were found to have a shared strain with LOCAL, while 50% of species with ≥100 unique NCBI strains had a shared strain with LOCAL. Alternatively, perhaps the decaying tail of pairwise kmer overlap of *E. coli* (Fig. S1C and S1D) is a signature suggestive of an organism whose recombination and mutation rates are too fast to have a finite population size. Increased numbers of sequenced Bacteroidetes isolates in the coming years will reveal if a similar decay occurs for strains of species in this phylum.

The identification of finite bacterial strain populations suggests that for some species we might be able to approach a complete sequencing of all strains. This sequencing effort combined with strain tracking algorithms to identify the frequency of each strain in shallow metagenomics datasets^21^ from tens of thousands of individuals could facilitate the association of specific bacterial strains with human health and disease to complement gene-based associations. Since the most frequent strains will likely be isolated first, this initiative would enable association of health outcomes with the most prevalent human associated microbes and enable studies to understand factors driving strain prevalence in the human population.

## Methods

### Bacterial genomes

Bacteria were isolated as previously described^14,36^. All bacterial genomes from the LOCAL cohort were sequenced with an Illumina HiSeq2500 or HiSeq4000. The NCBI RefSeq 156,403 bacterial genomes set was downloaded on May 27, 2019 using filters to exclude: partial genomes, derived environmental sources, derived metagenome, derived from single cell, genome length too large, genome length too small, high contig L50, low contig N50, low quality sequence, many frameshifted proteins, and anomalous.

### Quantifying kmer distances and average nucleotide identity (ANI)

The kmer overlap between any two genomes A and B was determined by generating a hash for genome A with kmer size 20 and quantifying the proportion of kmers shared in both genomes A and B divided by the total number of kmers in A. These distances were independently calculated in both directions. Given the focus of this manuscript on species and strain-level distances, particularly those of kmer overlap >0.98, we initially calculated the kmer overlap for the first 50,000 kmers in each genome and only performed the full genome comparison when this initial kmer coverage was >0.1. ANI was calculated using the fastANI algorithm^5^.

### Estimation of bacterial strain population sizes for each species

The frequency distribution of strain genomes that were found once, twice, etc.. for a given species as f_1_, f_2_ … f_N_ was determined by quantifying all pairwise genome kmer distances between all 156,403 genomes in the NCBI cohort. Given the large number of pairwise distances, genomes were clustered at the strain-level with a greedy heuristic algorithm that joined a genome into the cluster if any other genome in the cluster had >0.98 kmer overlap. The frequencies of f_1_, f_2_,…, f_N_ were calculated as the number of clusters of size 1, 2,…, N respectively. Population sizes estimated from these frequencies were calculated using the iChao algorithm based on Hill statistics. Population sizes estimated with the Mark and Recapture approach were estimated with the Chapman estimator 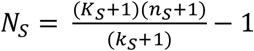 where *N_S_* is the number of strains in the population for species *S*, *n_S_* is the number of unique strains in the NCBI dataset for species S, *k_S_* is the number of species S strains in the LOCAL dataset that were also find in the NCBI database, and *K_s_* is the number of strains from species S in the LOCAL dataset.

### Data and code availability

Bacterial genomes for this study are available via NCBI BioProject PRJNA637878.

## Acknowledgments

This work was supported in part by the staff and resources of the Microbiome Translational Center and the Scientific Computing Division in Icahn School of Medicine at Mount Sinai. This work was supported by National Institutes of Health Grants (NIDDK DK112978 and NIDDK DK114133).

## Author contributions

J.J.F conceived the study and designed the experiments; E.S.S provided insights from infectious disease; I.M. developed the high throughput culturing and genome sequencing infrastructure. N.O.K., T.J.B., H.M., M.A.K., and S.P. collected Australian stool samples; I.M., A.C.L., and Z.L. isolated and sequenced the bacterial genomes; J.J.F., I.M., E.S.S. and V.A. analyzed data; J.J.F. wrote the manuscript. All authors read and approved the final manuscript.

---

**Figure S1.**
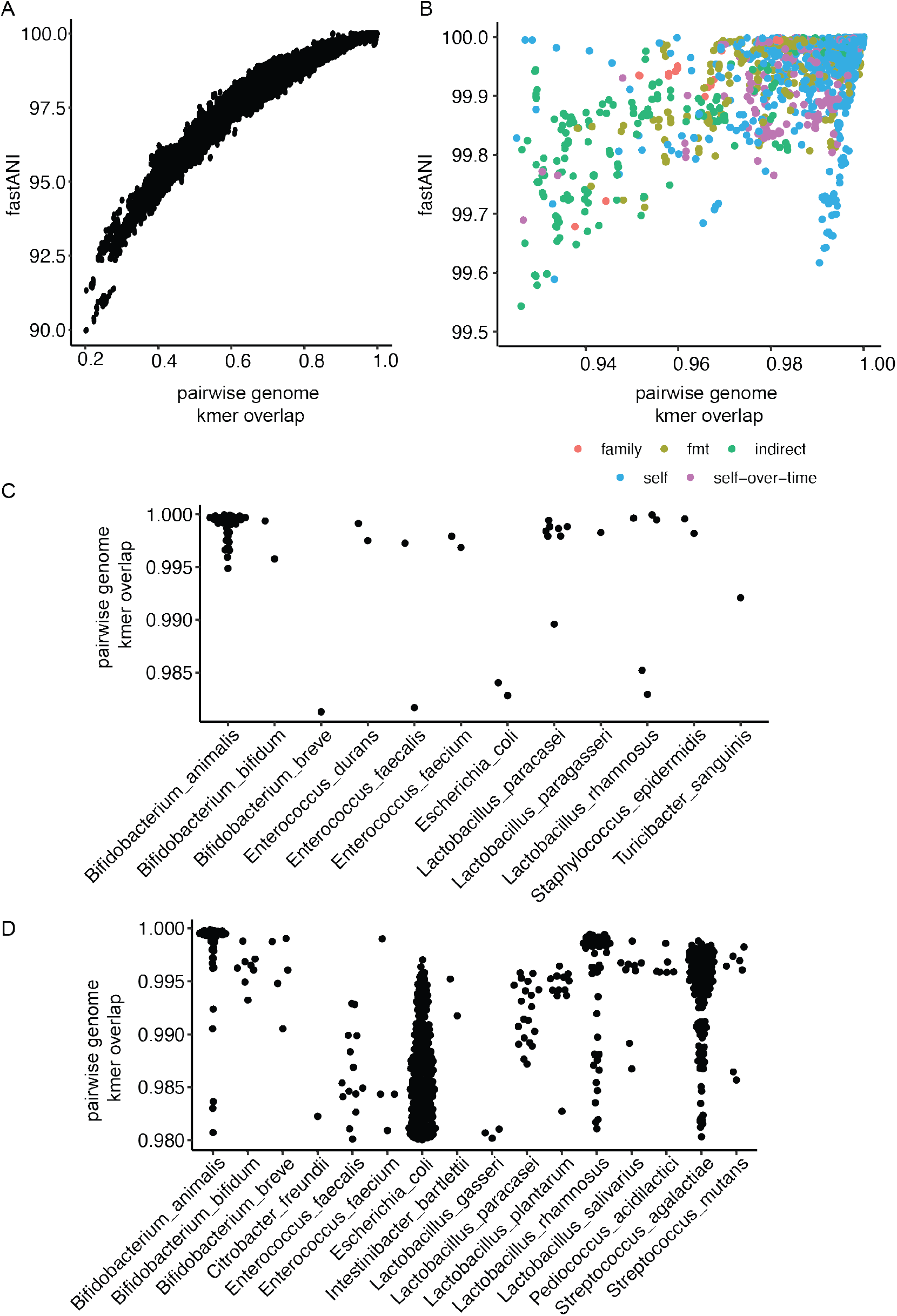
Comparing genome similarity metrics and observing the extreme tail of the kmer overlap distribution. **(A)** Overall, genome similarities measured with fastANI and kmer overlap are highly correlated with the fastANI metric appearing to approach saturation in resolving very similar genomes from the same species. **(B)** Empirically, the kmer overlap metric of >0.98 seems to delineate self-vs-self comparisons (magenta and blue points) more consistently than fastANI. **(C,D)** With few exceptions the kmer overlap comparisons >0.98 for each species are skewed towards kmer overlaps > 0.99.

